# Synaptic density imaging associates with performance but not self-report in autism

**DOI:** 10.64898/2026.10.05.755105

**Authors:** Adam J. Naples, Salih Cayir, Christine Cukar-Capizzi, Julie M. Wolf, Jenna E. McClelland, Emma C. Robertson, Shivani S. Sawant, David Matuskey, James C. McPartland

**Affiliations:** Center for Brain and Mind Health, Yale University, New Haven, CT; Child Study Center, Yale University, New Haven, CT; Department of Radiology and Biomedical Imaging, Yale University, New Haven, CT; Department of Psychiatry and Behavioral Sciences, McGovern Medical School, The University of Texas Health Science Center at Houston (UTHealth); Department of Psychiatry, Yale University, New Haven, CT; Department of Neurology, Yale University, New Haven, CT, United States

**Keywords:** Autism, Positron Emission Tomography, Synapse, SV2A

## Abstract

Autism is a common neurodevelopmental condition with considerable between-person variability and unknown biological etiology. Diagnosis of autism in adults relies on measurement of performance (e.g., clinician rating of social behavior, standardized assessment of cognition) and self-report of autism features. Our prior work revealed a strong relationship between clinician ratings of social function in autism and *in vivo* measures of synaptic density as measured by [^11^C]UCB-J PET (Matuskey et al., 2025). However, it is not understood how the measured aspects of the autism phenotype index underlying biology. Here, in a sample of 12 autistic adults, we address this question by relating synaptic density imaging to a range of performance measures, such as cognition and face recognition, in contrast to self-report measures. Our results identify a dissociation between these measurement types in that synaptic density correlates strongly with measures of performance but shows virtually no relationship with self-report. These findings suggest that, while self-report and performance measures share variance in characterizing phenotypes in autism, they diverge when mapping onto the underlying biology. Results suggest that higher synaptic density may index a general mechanism that facilitates social and cognitive performance across a range of tasks. These data also indicate that performance and self-report measures are not interchangeable in capturing brain-behavior correspondences in autism, a finding that can guide measure selection in future biomarker studies and clinical research.

## Introduction

Autism is a prevalent and heterogeneous neurodevelopmental condition with unclear biological underpinnings. Lack of insight into biological mechanisms has hindered progress in developing effective treatments and identifying objective biomarkers. The primary diagnostic criteria for autism remain behavioral differences in social interaction and communication. These differences are measured in clinical and research contexts using two measurement approaches: (1) direct measures of performance, e.g., clinician ratings of social behavior and cognitive testing; and (2) self- or caregiver reports of autistic features. While recent research has made progress in developing novel biomarkers,^1^ little is understood about the nature of clinical measures and their alignment with underlying biology.

Performance and self-report measures index different aspects of the autistic phenotype even as they span overlapping symptom domains such as social function.^2^ Self-report measures are cost-effective and practical for capturing variability across diverse symptom domains and contexts and uniquely measure subjective experience. However, self-report lacks an objective reference criterion, hindering comparability across individuals. For example, a question such as “Rate on a scale of one to five, ‘I have good self-confidence’” may yield the same numeric response from individuals whose internal states differ. In contrast, performance measures, such as cognitive tests or the Autism Diagnostic Observation Schedule (ADOS-2^3^), directly elicit behavior in real time, yielding more objective data that allows for direct comparisons between individuals and to external criteria. However, these measures are resource-intensive and require trained administrators. While both self-report and performance measures are used in biomarker research, it remains unknown which approach provides the most effective mapping, in a simple and direct way, to biological markers.

A recent advance in biomarker research is the development of positron emission tomography (PET) imaging of *in vivo* synaptic density.^4^ Genetic, *in vitro*, and post-mortem research has implicated changes at the synapse as a mechanism for autistic differences.^5^ Recently, we published the first study of *in vivo* synaptic density in autism and found lower synaptic density across all brain regions in autistic adults compared to non-autistic controls with [^11^C]UCB-J, although another group did not replicate these findings.^6,7^ Additionally, we found that, among autistic adults, synaptic density strongly correlated with clinician observation of social performance measured by the ADOS-2, such that lower synaptic density was associated with increased social-communicative difficulties.^6^ However, it is not known whether this relationship is specific to this measure of social performance or if it extends to other performance or self-report measures commonly used in autism biomarker research.

In this study, we expanded on this finding by investigating how performance measures (of both social and cognitive function) and self-report relate to synaptic density. In the same cohort of 12 autistic adults reported in Matuskey et al., 2025,^6^ we quantified the relationship between regional and global synaptic density (measured using PET imaging) and measures of performance (e.g., cognitive ability, face recognition, and emotion recognition) compared to self-report of autistic features. We hypothesized that performance measures, regardless of content area, would show a similarly strong relationship with synaptic density, while self-report measures would show weaker correlations.

## Results

Irrespective of content focus, performance measures were strongly associated with synaptic density in all brain regions, such that higher synaptic density was associated with higher levels of performance (Figure 1). There were no significant associations between self-report measures and synaptic density.

**Figure 1.**
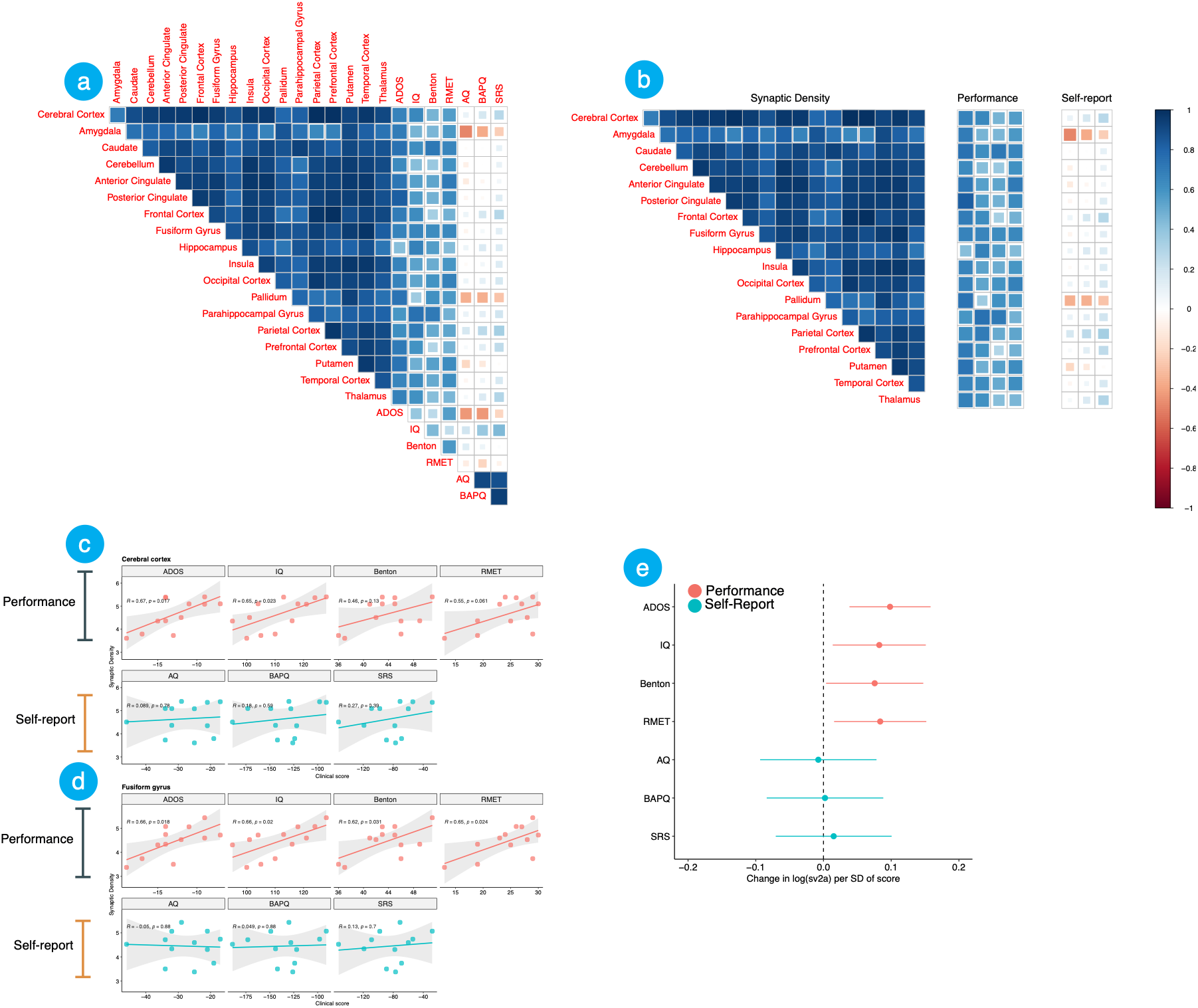
Associations between clinical measures and regional synaptic density. Figure 1a shows the correlation matrix between clinical measures and synaptic density. The size and color of the square indicate the magnitude and direction of the correlation. Figure 1b shows the matrix divided to highlight: (1) Correlations between different brain regions; (2) correlations between regional synaptic density and performance measures; (3) correlations between regional synaptic density and self-report measures. Performance measures show larger correlations with synaptic density than self-report measures, and the direction of these correlations is consistent across brain regions. Figures 1c and 1d show selected individual scatterplots between clinical measures and synaptic density in the entire cerebral cortex as an ROI (1c) and the fusiform gyrus (1d), a core node in the face processing network, which is implicated in social processing differences in autism.^18^ Figure 1e shows results of mixed-effects regressions with circles indicating fixed effects of the measure score in predicting synaptic density and horizontal lines indicating 95% confidence intervals. Performance measures show large effects with consistent directionality.

We quantified relationships between synaptic density and clinical measures in two ways. First, we calculated bivariate correlations between each clinical measure and synaptic density in each brain region. Excluding our previously reported relationship between the ADOS-2 and synaptic density, this effect was significant at *p* < .05 in 27 out of 54 comparisons with performance measures. Additionally, all non-significant correlations with performance measures were in the same direction and of similar magnitude, whereas there was no consistent pattern of relationships between synaptic density and self-report measures. We followed up each correlation with a permutation test to compare against the empirical null distribution, and patterns of significance were unchanged. All correlations are shown in Supplementary Table S1.

Second, to quantify the consistency of relationships across brain regions, we conducted separate mixed-effects regressions for each clinical measure. In each regression, synaptic density (log-transformed to account for between-region differences in mean and variance) was the dependent variable, and clinical score and brain region (excluding the overall cerebral cortex ROI) were fixed effects. For each performance measure, the fixed effect of the score was significant (IQ F = 6.82, *p* = .023; ADOS-2 F = 12.77, p = .004; Benton F = 5.27, p = .041; RMET F = 7.18, p = .02). No fixed effects were significant for self-report measures (AQ F = .04, *p* = .85; BAPQ F < 0.01, *p* = .956; SRS F = .15, *p* = .708). Full model results are in Table 1. Given the small sample size and exploratory nature of our approach, results were not corrected for multiple comparisons.

**Table 1.** Effects of score and region on synaptic density, by measure

| Measure | Score |  |  | Region |  |  |
| --- | --- | --- | --- | --- | --- | --- |
|  | *F* | *df* | *p* | *F* | *df* | *p* |
| Performance Measures |  |  |  |  |  |  |
| ADOS | 12.77 | 1, 12.0 | 0.004* | 368.64 | 16, 192.0 | <0.001 |
| IQ | 6.82 | 1, 12.0 | 0.023* | 368.64 | 16, 192.0 | <0.001 |
| Benton | 5.27 | 1, 12.0 | 0.041* | 368.64 | 16, 192.0 | <0.001 |
| RMET | 7.18 | 1, 12.0 | 0.020* | 368.64 | 16, 192.0 | <0.001 |
| Self-report Measures |  |  |  |  |  |  |
| AQ | 0.04 | 1, 12.0 | 0.850 | 368.64 | 16, 192.0 | <0.001 |
| BAPQ | 0.00 | 1, 12.0 | 0.956 | 368.64 | 16, 192.0 | <0.001 |
| SRS | 0.15 | 1, 12.0 | 0.708 | 368.64 | 16, 192.0 | <0.001 |

*Note.* ADOS, Autism Diagnostic Observation Schedule–2; RMET, Reading the Mind in the Eyes Test; IQ, Wechsler Abbreviated Scale of Intelligence–II; AQ, Autism Spectrum Quotient; BAPQ, Broad Autism Phenotype Questionnaire; SRS-A, Social Responsiveness Scale–Adult Self-Report. \* $p < 0.05$ .

Finally, we computed Pearson correlations between regional synaptic density (*BP*_ND_) and each clinical measure across all 87 FreeSurfer regions.The resulting brain maps illustrate the directionality and magnitude of association for each region (Figure 2), which was consistent with the primary ROI analyses. Overall, this pattern of results indicates that performance measures show consistent and large effect size relationships with synaptic density in autism.

**Figure 2.**
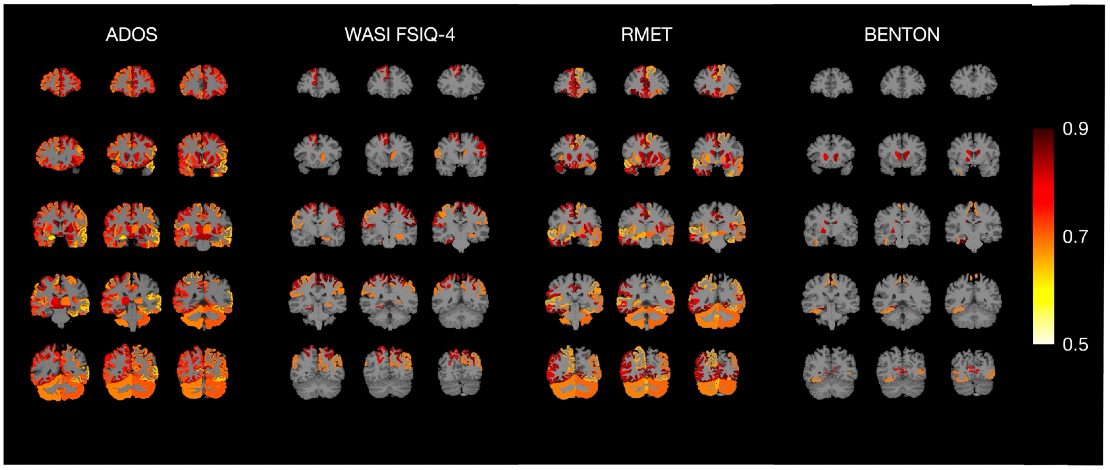
Correlation maps between regional *BP*_ND_ values and performance measures. Correlation maps were generated by assigning the calculated Pearson’s r value uniformly to all voxels within each corresponding FreeSurfer region and overlaying the resulting maps onto an MNI template T1-weighted MRI. The color scale is displayed only for regions reaching an uncorrected significance threshold of *p* < 0.05.

## Conclusion

In this study, performance, but not self-report, measures of autistic features correlated with synaptic density imaging, indicating that the same constructs captured with different methods relate differently to underlying biology. Moreover, performance measures tapping additional constructs of cognitive ability, face recognition, and emotion recognition were strongly related to synaptic density. This pattern of results suggests that synaptic density may broadly relate to cross-domain human cognitive and social performance in ways that are not reflected by self-report.

There are several potential explanations for the dissociation between self-report and performance measures. First, synaptic density and performance measures reflect contemporaneous measurement, i.e., they are measured in a constrained period of real time. In contrast, self-report measures are retrospective and represent the mental aggregation of experiences over time and across brain states. Second, many of these measures include items about how a person engages with others in social interactions, and thus depend on environmental factors (e.g., school or workplace) that may introduce error variance into the measures. Finally, as mentioned earlier, the absence of an external criterion to anchor responses across participants may limit the utility of self-report in making between-person comparisons. These hypotheses are supported by large-scale biomarker studies in autism using EEG and eye-tracking,^7,8^ which have found the strongest relationships with contemporaneous performance measures of IQ and face memory, and small or null relationships with caregiver-report measures.

Additionally, our results add to the literature linking cognitive ability and synaptic density by identifying, for the first time, a relationship between IQ and synaptic density in autism. This relationship may reflect increased computational efficiency across the entire brain driven by synaptic connectivity and partially explain our findings in the emotion and face recognition tasks, which place demands on working memory and vocabulary in addition to social perception. Similar relationships between cognitive performance, e.g., working memory and IQ, and synaptic density, have been reported across many conditions.^9^ Of note, synaptic density in the fusiform gyrus, an area that subserves face processing, showed the strongest correlations with the emotion and face recognition tasks. However, these correlations were not limited to the fusiform and were similar across the whole brain. This result suggests that while performance on these social-cognitive tasks may benefit from region-specific variation, it does not appear to be specific to these regions.

This study has several limitations that necessitate further research. First, our sample only included autistic adults without co-occurring intellectual disability. This limits our ability to generalize these findings to non-autistic people, to autistic people with intellectual disability, or to younger cohorts. Second, our small sample size reflects the practical limitations of PET research and lacks power to detect small effects. Third, some participants were taking psychiatric medications, although Sv2a medications (e.g., levetiracetam) were excluded, we cannot rule out the possibility of medication effects.^10^ Finally, while our results show a dissociation between types of measures, we cannot know whether this finding generalizes to all self-report measures or is specific to those tools used in this sample. Further research is needed to determine which facets of measurement (e.g., contemporaneous administration or external criterion) explain these results.

In conclusion, these results expand our understanding of how the molecular mechanisms of autism relate to distinct forms of phenotypic measurement. Specifically, our results suggest that performance measurement approaches may be most fruitful for identifying brain-behavior relationships. This is germane to the design of future biomarker studies in autism.

## Materials and Methods

Autistic participants (n = 12; mean (SD) age 25 (4) years; six males) met gold-standard research criteria for ASD diagnosis (ADOS-2/Brief Observation of Symptoms of Autism [BOSA; administered to accommodate pandemic-induced masking requirements] and DSM-5 clinical diagnosis).^3,11^ The primary metric of social-communicative performance was clinician rating on the ADOS-2 Module 4 and the BOSA version F2/Module 4. Data were combined across these two measures as reported previously.^6^ Self-report measures included the Social Responsiveness Scale – Adult Self-Report, Second Edition [SRS]^12^; Autism-Spectrum Quotient [AQ]^13^; Broad Autism Phenotype Questionnaire [BAPQ]^14^). Performance measures included the Benton Facial Recognition task [Benton]^15^; The “Reading the Mind in the Eyes” Task [RMET].^16^ Cognitive testing used the Wechsler Abbreviated Scale of Intelligence – Second Edition (WASI-II)^17^. To aid in interpretation of directionality of results across measures, measures of autistic features (ADOS,SRS,BAPQ,AQ) were reverse scored. In this way, higher scores on all measures reflected better performance or reduced autistic features. All participants underwent a single MRI and dynamic [^11^C]UCB-J PET scan on a High-Resolution Research Tomograph (Siemens). The outcome measure was non-displaceable binding potential (*BP*_ND_) with the centrum semiovale as a reference region. Full acquisition, reconstruction, and quantification details are provided in the Supplementary Methods and reported previously.^6^

Eligibility was confirmed through comprehensive psychiatric and medical histories, physical examination, and laboratory assessments. The study was performed under a protocol approved by the Yale University Human Investigation Committee and the Yale New Haven Hospital Radiation Safety Committee. Written informed consent was obtained from all participants after a complete explanation of study procedures, including clarification that discontinuation of participation would not yield negative consequences.

## Acknowledgements

This research was supported by NARSAD (Matuskey), NIMH R21 MH113955 (Matuskey, McPartland) and the Sebastian Innovation Fund in Autism (Naples, Matuskey). We thank the staff of the Yale PET Center and the Yale Magnetic Resonance Research Center (MRRC) for their help in conducting this work. Finally, we would like to thank all of our participants for helping to generate new knowledge and for making an important contribution to autism research.

