## Supplement for "Synaptic density imaging associates with performance but not self-report in autism"

SUPPLEMENTARY MATERIALS

**Table S1. Correlations between synaptic density and clinical measures in all regions for participants with autism. *p*_perm indicates permutation calculated *p*-values. Cells highlighted in red indicate *p* < .05.**

| Measure | Region | r | *p* | *p*_perm |
|---|---|---|---|---|
| **Performance Measures** | | | | |
| ADOS | Cerebral Cortex | 0.67 | 0.017 | 0.016 |
| ADOS | Amygdala | 0.70 | 0.012 | 0.011 |
| ADOS | Caudate | 0.75 | 0.005 | 0.007 |
| ADOS | Cerebellum | 0.63 | 0.027 | 0.024 |
| ADOS | Anterior Cingulate | 0.72 | 0.008 | 0.009 |
| ADOS | Posterior Cingulate | 0.82 | 0.001 | 0.001 |
| ADOS | Frontal Cortex | 0.69 | 0.013 | 0.012 |
| ADOS | Fusiform Gyrus | 0.66 | 0.018 | 0.017 |
| ADOS | Hippocampus | 0.42 | 0.169 | 0.169 |
| ADOS | Insula | 0.66 | 0.019 | 0.021 |
| ADOS | Occipital Cortex | 0.66 | 0.019 | 0.025 |
| ADOS | Pallidum | 0.74 | 0.006 | 0.008 |
| ADOS | Parahippocampal Gyrus | 0.58 | 0.047 | 0.048 |
| ADOS | Parietal Cortex | 0.61 | 0.034 | 0.036 |
| ADOS | Prefrontal Cortex | 0.71 | 0.010 | 0.010 |
| ADOS | Putamen | 0.74 | 0.006 | 0.009 |
| ADOS | Temporal Cortex | 0.66 | 0.020 | 0.018 |
| ADOS | Thalamus | 0.68 | 0.016 | 0.016 |
| IQ | Cerebral Cortex | 0.65 | 0.023 | 0.025 |
| IQ | Amygdala | 0.47 | 0.127 | 0.121 |
| IQ | Caudate | 0.62 | 0.033 | 0.038 |
| IQ | Cerebellum | 0.43 | 0.165 | 0.157 |
| IQ | Anterior Cingulate | 0.55 | 0.064 | 0.067 |
| IQ | Posterior Cingulate | 0.56 | 0.057 | 0.052 |
| IQ | Frontal Cortex | 0.63 | 0.029 | 0.031 |
| IQ | Fusiform Gyrus | 0.66 | 0.020 | 0.023 |
| IQ | Hippocampus | 0.68 | 0.016 | 0.016 |
| IQ | Insula | 0.53 | 0.076 | 0.080 |
| IQ | Occipital Cortex | 0.52 | 0.081 | 0.084 |
| IQ | Pallidum | 0.35 | 0.263 | 0.261 |
| IQ | Parahippocampal Gyrus | 0.74 | 0.006 | 0.006 |
| IQ | Parietal Cortex | 0.69 | 0.013 | 0.016 |
| IQ | Prefrontal Cortex | 0.60 | 0.040 | 0.044 |
| IQ | Putamen | 0.49 | 0.107 | 0.108 |
| IQ | Temporal Cortex | 0.62 | 0.031 | 0.036 |
| IQ | Thalamus | 0.60 | 0.039 | 0.043 |
| Benton | Cerebral Cortex | 0.46 | 0.129 | 0.130 |
| Benton | Amygdala | 0.45 | 0.138 | 0.143 |

| | | | | |
|---|---|---|---|---|
| Benton | Caudate | 0.72 | 0.009 | 0.009 |
| Benton | Cerebellum | 0.38 | 0.224 | 0.224 |
| Benton | Anterior Cingulate | 0.52 | 0.080 | 0.084 |
| Benton | Posterior Cingulate | 0.47 | 0.120 | 0.128 |
| Benton | Frontal Cortex | 0.34 | 0.283 | 0.292 |
| Benton | Fusiform Gyrus | 0.62 | 0.031 | 0.028 |
| Benton | Hippocampus | 0.55 | 0.061 | 0.058 |
| Benton | Insula | 0.49 | 0.105 | 0.105 |
| Benton | Occipital Cortex | 0.59 | 0.042 | 0.038 |
| Benton | Pallidum | 0.61 | 0.036 | 0.036 |
| Benton | Parahippocampal Gyrus | 0.70 | 0.011 | 0.011 |
| Benton | Parietal Cortex | 0.50 | 0.096 | 0.095 |
| Benton | Prefrontal Cortex | 0.30 | 0.347 | 0.344 |
| Benton | Putamen | 0.54 | 0.068 | 0.064 |
| Benton | Temporal Cortex | 0.50 | 0.095 | 0.098 |
| Benton | Thalamus | 0.43 | 0.167 | 0.169 |
| RMET | Cerebral Cortex | 0.55 | 0.061 | 0.061 |
| RMET | Amygdala | 0.57 | 0.051 | 0.050 |
| RMET | Caudate | 0.65 | 0.022 | 0.022 |
| RMET | Cerebellum | 0.60 | 0.039 | 0.041 |
| RMET | Anterior Cingulate | 0.71 | 0.010 | 0.009 |
| RMET | Posterior Cingulate | 0.63 | 0.028 | 0.024 |
| RMET | Frontal Cortex | 0.47 | 0.121 | 0.116 |
| RMET | Fusiform Gyrus | 0.65 | 0.024 | 0.022 |
| RMET | Hippocampus | 0.43 | 0.159 | 0.160 |
| RMET | Insula | 0.59 | 0.044 | 0.046 |
| RMET | Occipital Cortex | 0.68 | 0.015 | 0.014 |
| RMET | Pallidum | 0.58 | 0.050 | 0.045 |
| RMET | Parahippocampal Gyrus | 0.47 | 0.119 | 0.131 |
| RMET | Parietal Cortex | 0.52 | 0.082 | 0.082 |
| RMET | Prefrontal Cortex | 0.48 | 0.115 | 0.120 |
| RMET | Putamen | 0.63 | 0.029 | 0.028 |
| RMET | Temporal Cortex | 0.62 | 0.033 | 0.034 |
| RMET | Thalamus | 0.45 | 0.142 | 0.146 |
| **Self-report Measures** | | | | |
| AQ | Cerebral Cortex | 0.09 | 0.783 | 0.775 |
| AQ | Amygdala | -0.49 | 0.108 | 0.110 |
| AQ | Caudate | -0.01 | 0.973 | 0.971 |
| AQ | Cerebellum | -0.08 | 0.811 | 0.817 |
| AQ | Anterior Cingulate | -0.10 | 0.754 | 0.756 |
| AQ | Posterior Cingulate | -0.05 | 0.884 | 0.883 |
| AQ | Frontal Cortex | 0.08 | 0.812 | 0.822 |

| | | | | |
|---|---|---|---|---|
| AQ | Fusiform Gyrus | -0.05 | 0.877 | 0.880 |
| AQ | Hippocampus | 0.07 | 0.821 | 0.829 |
| AQ | Insula | -0.02 | 0.940 | 0.938 |
| AQ | Occipital Cortex | 0.07 | 0.825 | 0.820 |
| AQ | Pallidum | -0.36 | 0.257 | 0.259 |
| AQ | Parahippocampal Gyrus | 0.08 | 0.798 | 0.803 |
| AQ | Parietal Cortex | 0.17 | 0.589 | 0.588 |
| AQ | Prefrontal Cortex | 0.04 | 0.911 | 0.912 |
| AQ | Putamen | -0.20 | 0.531 | 0.535 |
| AQ | Temporal Cortex | -0.04 | 0.893 | 0.895 |
| AQ | Thalamus | 0.04 | 0.900 | 0.902 |
| BAPQ | Cerebral Cortex | 0.18 | 0.586 | 0.594 |
| BAPQ | Amygdala | -0.37 | 0.231 | 0.225 |
| BAPQ | Caudate | 0.01 | 0.986 | 0.986 |
| BAPQ | Cerebellum | 0.00 | 1.000 | 1.000 |
| BAPQ | Anterior Cingulate | -0.04 | 0.901 | 0.907 |
| BAPQ | Posterior Cingulate | -0.03 | 0.939 | 0.940 |
| BAPQ | Frontal Cortex | 0.17 | 0.595 | 0.603 |
| BAPQ | Fusiform Gyrus | 0.05 | 0.880 | 0.887 |
| BAPQ | Hippocampus | 0.15 | 0.633 | 0.630 |
| BAPQ | Insula | 0.05 | 0.874 | 0.880 |
| BAPQ | Occipital Cortex | 0.10 | 0.746 | 0.746 |
| BAPQ | Pallidum | -0.35 | 0.265 | 0.270 |
| BAPQ | Parahippocampal Gyrus | 0.11 | 0.724 | 0.721 |
| BAPQ | Parietal Cortex | 0.26 | 0.410 | 0.422 |
| BAPQ | Prefrontal Cortex | 0.13 | 0.694 | 0.690 |
| BAPQ | Putamen | -0.12 | 0.713 | 0.713 |
| BAPQ | Temporal Cortex | 0.07 | 0.842 | 0.838 |
| BAPQ | Thalamus | 0.15 | 0.631 | 0.632 |
| SRS | Cerebral Cortex | 0.27 | 0.390 | 0.395 |
| SRS | Amygdala | -0.25 | 0.442 | 0.454 |
| SRS | Caudate | 0.10 | 0.753 | 0.756 |
| SRS | Cerebellum | 0.09 | 0.772 | 0.771 |
| SRS | Anterior Cingulate | 0.05 | 0.869 | 0.876 |
| SRS | Posterior Cingulate | 0.14 | 0.673 | 0.673 |
| SRS | Frontal Cortex | 0.29 | 0.362 | 0.366 |
| SRS | Fusiform Gyrus | 0.13 | 0.699 | 0.685 |
| SRS | Hippocampus | 0.15 | 0.653 | 0.661 |
| SRS | Insula | 0.14 | 0.654 | 0.656 |
| SRS | Occipital Cortex | 0.17 | 0.595 | 0.589 |
| SRS | Pallidum | -0.28 | 0.372 | 0.370 |
| SRS | Parahippocampal Gyrus | 0.19 | 0.546 | 0.551 |

| | | | | |
|---|---|---|---|---|
| SRS | Parietal Cortex | 0.34 | 0.284 | 0.281 |
| SRS | Prefrontal Cortex | 0.26 | 0.420 | 0.424 |
| SRS | Putamen | -0.01 | 0.965 | 0.966 |
| SRS | Temporal Cortex | 0.17 | 0.605 | 0.622 |
| SRS | Thalamus | 0.29 | 0.353 | 0.367 |

Pearson's *r* values and associated *P*-values (uncorrected for multiple comparisons) and empirical P-values are shown for each regional correlation analysis. ADOS, Autism Diagnostic Observation Schedule–2; RMET, Reading the Mind in the Eyes Test; IQ, Wechsler Abbreviated Scale of Intelligence–II; AQ, Autism Spectrum Quotient; BAPQ, Broad Autism Phenotype Questionnaire; SRS, Social Responsiveness Scale–Adult Self-Report.

### PET METHODS

#### Positron Emission Tomography

Each participant underwent a [$^{11}$C]UCB-J PET scan, lasting up to 90 minutes, using the High-Resolution Research Tomograph (HRRT, Siemens CTI, Knoxville, TN, USA). The [$^{11}$C]UCB-J radiotracer was synthesized on-site using established methods[1] and administered intravenously as a bolus via an automated infusion pump (Harvard PHD 22/2000, Harvard Apparatus). Prior to tracer injection, a 6-minute transmission scan was performed for attenuation correction. Head motion was continuously monitored using the Polaris Vicra optical tracking system (NDI Systems, Waterloo, Ontario, Canada). The collected data were reconstructed with corrections for attenuation, normalization, scatter, random coincidences, dead time, and motion using the MOLAR algorithm.[2] Previous studies have demonstrated that the primary outcome measure, binding potential ($BP_{ND}$), can be reliably calculated without arterial blood sampling in participants with autism,[3] Parkinson's disease,[4] and Alzheimer's disease.[5]

#### Quantitative analysis of PET data

Summed PET images were registered to the subject's T1-weighted MR images and then registered to an MR template. The primary outcome measure was $BP_{ND}$, calculated using the simplified reference tissue model (SRTM2).[6] Parametric images of $BP_{ND}$ were generated using the centrum semiovale as a reference region and a fixed $k_2'$ value (clearance rate constant, $k_2$, of the reference region). The centrum semiovale was defined using an average of individual white matter probability maps optimized to represent the core of the centrum semiovale.[7] The regions of interest (ROIs) were taken from the FreeSurfer 6.0 atlas (based on the Desikan-Killiany atlas) and were partial volume corrected (PVC) using the Iterative Yang algorithm.[8,9] The fixed $k_2'$ was a population-averaged value for the centrum semiovale derived from previous one-tissue compartment model analyses with arterial input (0.031 ± 0.005 min$^{-1}$, n = 17).[7] Based on prior autism neuroimaging literature, the primary ROI for PET analyses was the whole cerebral cortex. Seventeen exploratory ROIs were examined: prefrontal cortex, frontal cortex, anterior cingulate cortex, posterior cingulate cortex, parietal cortex, temporal cortex, occipital cortex, insula, fusiform gyrus, parahippocampal cortex, hippocampus, amygdala, caudate, putamen, pallidum, thalamus, and cerebellum cortex. Left and right hemisphere values were averaged. As a complementary analysis, Pearson correlations were computed between $BP_{\mathrm{ND}}$ and each clinical measure across all 87 FreeSurfer regions (68 cortical, 18 subcortical, and the vermis). For visualization, each region's r value was assigned uniformly to all voxels within that region and overlaid on an MNI template T1-weighted image (Figure 2). These analyses were exploratory and uncorrected for multiple comparisons.
